# Leveraging microtopography to pattern multi-oriented muscle actuators

**DOI:** 10.1101/2024.07.31.606059

**Authors:** Tamara Rossy, Laura Schwendeman, Maheera Bawa, Pavankumar Umashankar, Ritu Raman

**Affiliations:** Department of Mechanical Engineering, Massachusetts Institute of Technology, Cambridge, MA, USA 02139

## Abstract

Engineering skeletal muscle tissue with precisely defined alignment is of significant importance for applications ranging from drug screening to biohybrid robotics. Aligning 2D contractile muscle monolayers, which are compatible with high-content imaging and can be deployed in planar soft robots, typically require micropatterned cues. However, current protocols for integrating microscale topographical features in extracellular matrix hydrogels require expensive microfabrication equipment and multi-step procedures involving error-prone manual handling steps. To address this challenge, we present STAMP (Simple Templating of Actuators via Micro-topographical Patterning), an easily accessible and cost-effective one-step method to pattern microtopography of various sizes and configurations on the surface of hydrogels using reusable 3D printed stamps. We demonstrate that STAMP enables precisely controlling the alignment of mouse and human skeletal muscle fibers, and thus their force-generating axes, without impacting their maturation or function. To showcase the versatility of our technique, we designed a planar soft robot inspired by the iris, which leverages spatially segregated regions of concentric and radial muscle fibers to control pupil dilation. Optogenetic skeletal muscle fibers grown on a STAMPed iris substrates formed a multi-oriented actuator, and selective light stimulation of the radial and concentric fibers was used to control the function of the iris, including pupil constriction. Computational modeling of the biohybrid robot as an active bilayer matched experimental outcomes, showcase the robustness of our method of designing, fabricating, and testing planar biohybrid robots capable of complex multi-degree-of-freedom motion.

## Introduction

Skeletal muscle is a key component of the locomotor system, enabling voluntary movement through force generation.^1,2^ Engineered muscle tissues capable of producing measurable forces are valuable *in vitro* tools for screening new therapeutic strategies for neuromuscular pathologies.^3–7^ Engineered skeletal muscle has also emerged as an excellent candidate actuator for soft “biohybrid” robotics, owing to its energy efficiency, adaptability, and the ability to precisely trigger contraction via electrical or optical stimulation.^8–10^ An important consideration for all these applications is precisely patterning the alignment of multinucleated muscle fibers within engineered tissues.^11,12^

In 3D engineered muscles, fiber alignment is typically achieved by tensioning the tissue between two posts, yielding global alignment of fibers parallel to the axis of imposed tension.^13,14^ Deflection of the posts in response to muscle contraction has proven a useful method of monitoring tissue function in physiological and pathological states, as well as for converting tissue forces into useful robotic functions, such as walking and gripping.^15–18^ With the rise of high-content imaging, engineered contractile 2D muscle monolayers have been of increasing interest in the context of drug screening, as they enable rapid and scalable monitoring of tissue morphology and function.^5,7,19,20^ Precision alignment and compatibility with live cell imaging is particularly important in this application, as random alignment of muscle fibers may result in contractile forces and dynamics that are not directly comparable between biological replicates, thus rendering it difficult to make clear distinctions between healthy, diseased, and treated tissues. Patterning contractile 2D skeletal muscle is also interesting for biohybrid robotics, as it would enable deploying biological actuators in planar formats that enable multi-degree-of-freedom motions such as twisting, coiling, and swimming that have been accomplished with 2D cardiac muscle.^21–25^ However, accomplishing this goal requires the ability to pattern skeletal muscle monolayers that replicate the complex multi-oriented alignment geometries observed in native tissue, such as multipennate and circular muscles, and can be sustained in culture over several weeks.

For many years, 2D skeletal muscle monolayers were not considered useful or robust for real-world applications, as they delaminate from rigid substrates within a few days of differentiation, thus preventing long-term monitoring of morphology and contractile function.^26,27^ Early efforts to pattern adhesion-promoting proteins such as fibronectin, gelatin, and Matrigel on stiff plastic/glass substrates using microfabricated stamps thus proved successful at aligning skeletal muscle fibers over a few days in culture,^26,28^ but were not suitable for maintaining contractile monolayers over several weeks or months. As an alternative, several studies reported that muscle monolayers could be maintained for longer time periods on rigid microfabricated scaffolds coated with adhesion-promoting biopolymers if the scaffolds contained 3D microtopography, i.e. aligned grooves. While effective, these methods typically rely on complex multi-step protocols that require microfabrication facilities, and also use opaque materials such as silicon wafers as 3D scaffold materials which impairs live cell microscopy.^29–31^ Moreover, such rigid materials are not suitable scaffolds for planar soft robots which typically require compliant biocompatible polymers such as poly (dimethyl siloxane) and gelatin methacrylate to demonstrate large deformations in response to muscle actuation.^22,32^

Approaches to align 2D muscle monolayers by patterning grooves directly in soft extracellular-matrix mimicking hydrogels such as fibrin or collagen would overcome imaging limitations, and also better recapitulate the soft and viscoelastic mechanical properties of native tissue.^33^ However, hydrogel micro-molding frequently requires tedious fabrication protocols comprising error-prone manual handling steps (e.g. manually transferring and flipping over a layer of aligned muscle into a new culture dish^34^), or access to microfabrication equipment that are limited to photosensitive polymers,^35^ which limits the range of materials that can be used as well as the accessibility of the technique.

To address these issues, we developed a cost-effective, readily accessible, and one-step extracellular matrix molding approach to generate sheets of precisely aligned, contractile muscle in a variety of culture formats and from different cell sources. Our robust and reproducible method, termed “Simple Templating of Actuators via Micro-topographical Patterning” (STAMP), uses 3D printed stamps to patterning microscopic ridges into natural hydrogels cast in any format, including standard multi-well plates. We have designed and optimized a bubble-free gel-loading and defect-limiting mold release strategy to enable high-fidelity patterning of micro-topographical cues. Moreover, we show that a simple ultrasonication cleaning strategy can be deployed to enable sustainable re-use of stamps over multiple cycles.

To understand how cell size and origin impacts muscle alignment efficiency, fiber morphology, and contractile functionality, we seeded skeletal myoblasts from mouse and human origin on STAMPed fibrin hydrogels with varying groove geometries ranging from 1-10x the size of single cells in suspension. We found that microgrooves enabled controlling the orientation of muscle fibers irrespective of cell source and demonstrated similar patterning efficiency across a wide range of groove sizes, likely by constraining the direction of growth and fusion of myoblasts. Grooved and non-patterned engineered tissues contracted with comparable forces in response to electrical stimulation, indicating similar maturity of muscle fibers. Interestingly, substrate microtopography did not significantly influence the width or fusion index of individual muscle fibers, showing that our method allows independently manipulating the spatial configuration of muscle fibers without compromising tissue functionality.

As a showcase of the versatility of our method, we leveraged STAMP to pattern complex multi-degree-of-freedom motion in a planar muscle actuator. Given the difficulty of patterning muscle fiber alignment in multi-oriented geometries, biohybrid robots have thus far relied on proof-of-concept demonstrations of robots that walk, grip, or swim in response to one degree-of-freedom (DOF) motion from unidirectionally aligned 3D muscle tissues.^16–18,35,36^ The ability to simply, efficiently, and precisely pattern the alignment of contractile muscle fibers in any spatial geometry, while retaining their force-generation capacity, opens up the possibility of designing *multi*-oriented architectures capable of *multi*-DOF motion.

We fabricated a fibrin layer patterned with a series of concentric and radial microgrooves arranged around a circular hole, mimicking the native architecture of the iris muscles that control pupil dilation in the eye. Computational modeling of muscle monolayers on fibrin substrates, informed by our experiments comparing the effect of groove size and cell source, enabled predictive design of an iris-mimic that could control dilation of a “pupil” in response to triggerable muscle contraction. Optogenetic mouse muscle cells seeded on this substrate were differentiated in a multi-DOF muscle configuration. Controlling the contraction of the concentric and radial layers separately by selectively shining blue light on either region showcased regional specificity in contraction direction, and stimulation of the iris was used to control pupil constriction. Experimental results matched computational predictions, indicating significant future potential in designing and deploying planar biohybrid robots capable of complex multi-DOF motion. Beyond robotics, we anticipate STAMP will provide the tissue engineering community with a rapid, widely accessible, cost-efficient, and versatile way of introducing microscale topographical cues in ECM hydrogels without the need for complex multi-step microfabrication methodologies.

## Results

### Simple Templating of Actuators via Micro-topographical Patterning (STAMP)

To generate aligned muscle tissues in a fast and versatile manner, and explore the impact of groove size and cell source on tissue morphology and function, we developed a one-step hydrogel molding protocol that can be applied to a variety of cell culture formats, such as multi-well plates and microfluidic chips as well as a range of hydrogel crosslinking chemistries. Specifically, we 3D printed stamps that fit in commercially available 24-well plates and used them to pattern vertically aligned (90°) microgrooves into hydrogels cast in each well. Stamps were attached to a holder with an integrated “bubble release” feature to prevent formation of defects commonly seen in molding viscous pre-crosslinked hydrogels (Figure 1a-c and Supplementary Figure 1). We printed the holders as pairs connected by a rigid beam (Figure 1b) to facilitate vertical alignment of microgrooves with respect to the multi-well plate. We UV-sterilized the stamps before immersing them in a solution containing 1% bovine serum albumin for 1 hour to facilitate subsequent mold release.^37^ Stamps were dried and briefly rinsed in ultrapure water prior to use.

**Figure 1:**
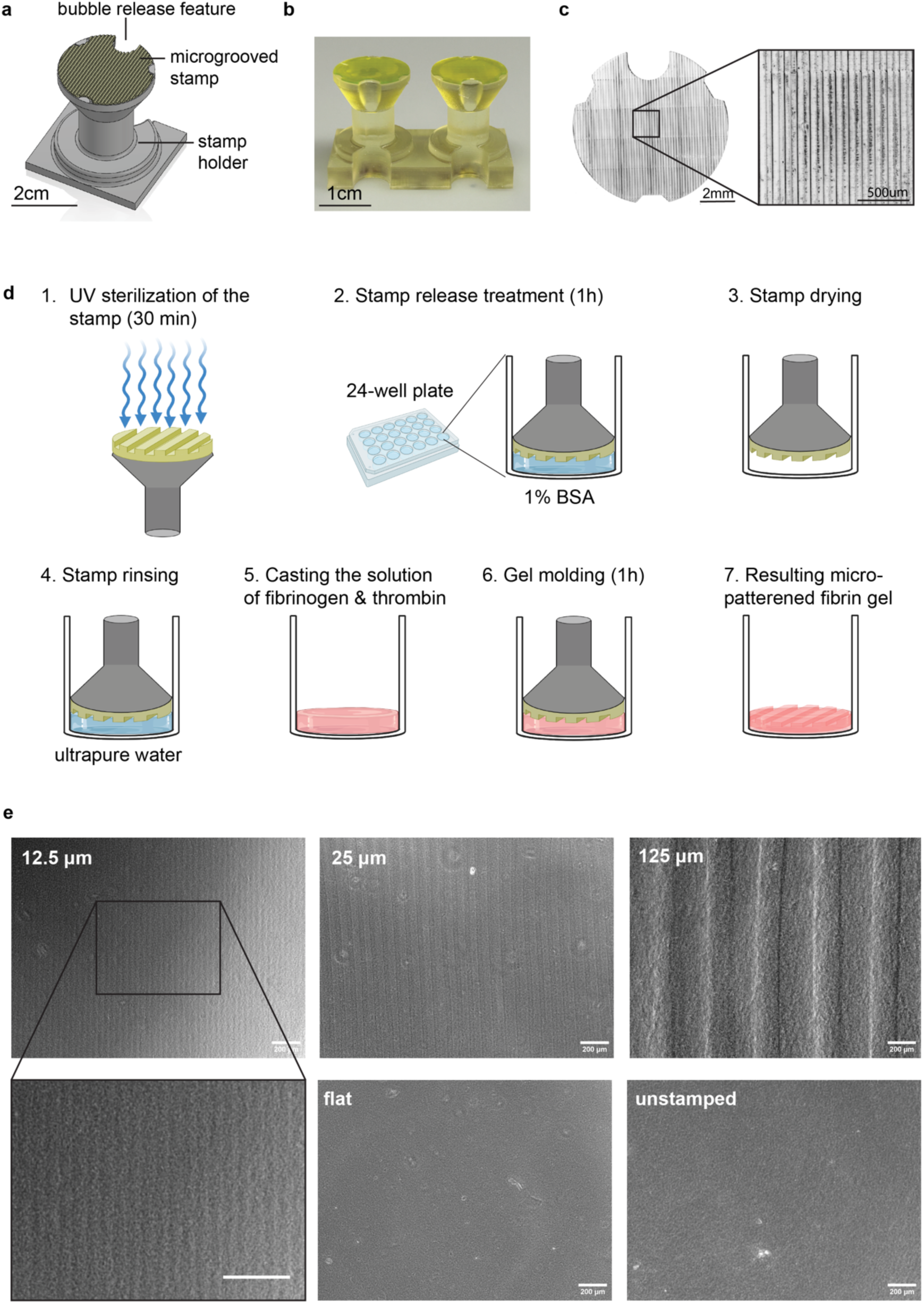
STAMP, a simple and versatile method to fabricate micro-grooved hydrogel substrates. **a.** Computer-assisted design (CAD) model of the micropatterned stamp and holder and **b-c.** resulting 3D-printed parts. **d.** Schematic steps of the stamp treatment and hydrogel patterning processes (created with Biorender.com). **e.** Brightfield microscopy pictures of fibrin gels harboring grooves of different sizes (equal width and depth), patterned with an non-grooved stamp (‘flat’), or not stamped at all.

To fabricated micro-grooved hydrogels, we cast a liquid solution of fibrinogen monomer (8 mg/mL) and thrombin crosslinker (100U/mL) into each well of a 24-well plate, given widespread use of fibrin as a substrate for skeletal muscle culture.^38–40^ Specifically, we followed our previously established protocols for generating fibrin with rheological properties that sustain longitudinal culture of skeletal muscle monolayers over several weeks.^41^ Immediately after casting the liquid prepolymer solution, we placed the stamps into the wells and incubated the setup for 1h at 37°C in a cell culture incubator to enable polymerization into a crosslinked fibrin hydrogel. Finally, we removed the stamps from the wells, showcasing efficient transfer of microgrooves onto the surface of the hydrogel (Figure 1d). The features of the pattern, such as the depth and width of the grooves can be easily varied by modifying the computer-assisted design (CAD) model of the stamp (Figure 1e). Furthermore, stamps can be reused after a quick cleaning protocol comprising rinsing, sonication and UV sterilization (Methods and Supplementary Figure 2). Overall, our method offers a rapid and versatile way to engineer the micro-scale topography of translucent and mechanically compliant cell culture substrates.

To explore whether and how the interplay between cell size and groove size influences muscle alignment, we printed stamps with groove dimensions (equal depth and width) of 1x, 2x, and 10x the size of single muscle cells of mouse (C2C12) and human (skMDC, Cook Myosite) origin (Figure 2a-b and Supplementary Figure 3), namely 12.5, 25 and 125 μm. We also printed a flat stamp devoid of microgrooves to verify that potential differences in cellular orientation were not a result of the gel stamping or mold-release coating process. Different variants of stamps were used to pattern fibrin gels as described above, in triplicates, prior to seeding with either mouse or human muscle cells (Figure 2a). As an additional control, we also seeded cells onto unstamped fibrin gels.

**Figure 2:**
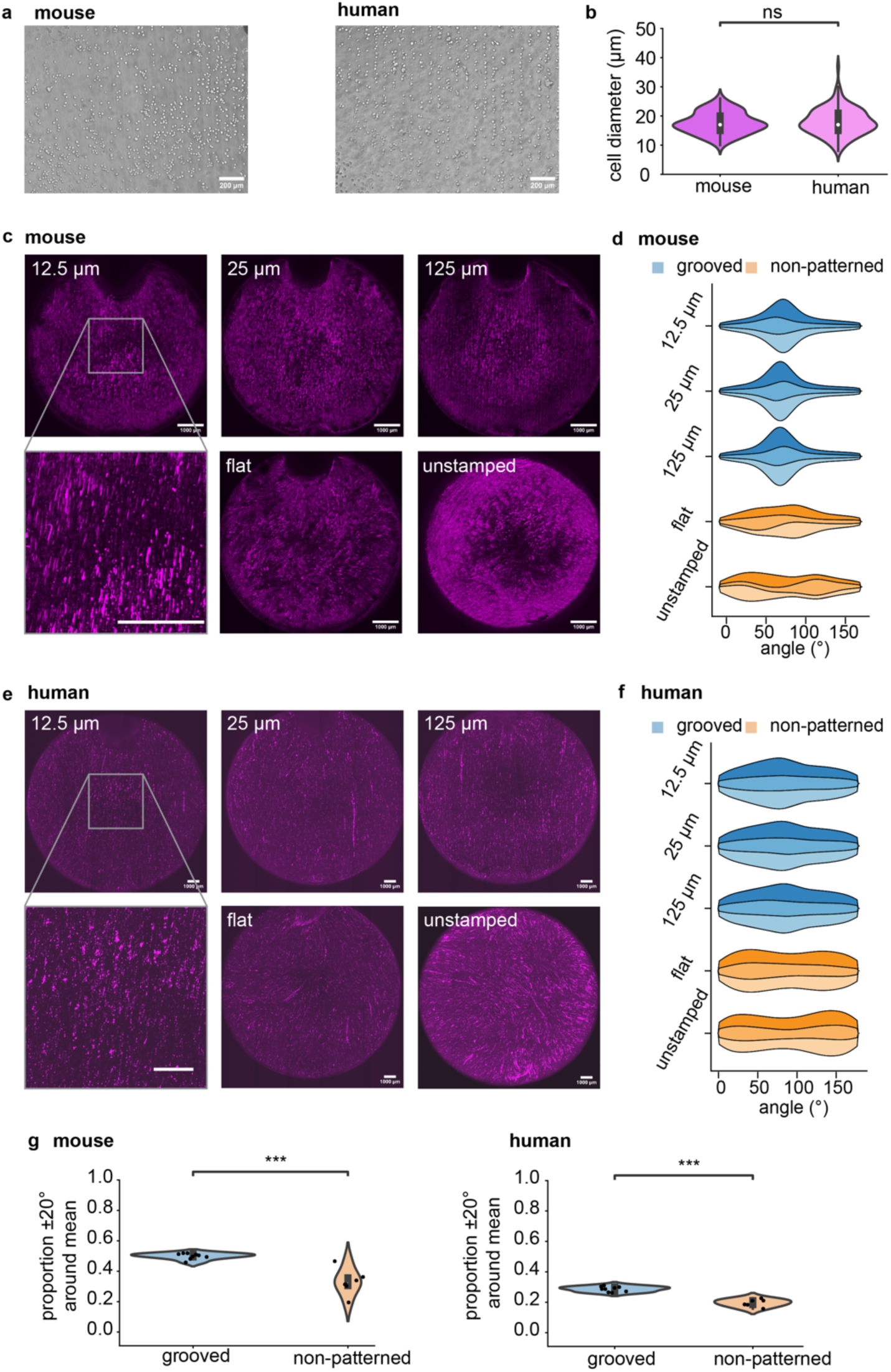
Effect of micro-grooved fibrin substrates on the orientation of muscle cells. **a.** Mouse and human myoblasts upon seeding on a fibrin scaffold containing 25 μm-wide grooves. Scale bar: 200 μm. **b.** Diameter of single mouse and human myoblasts in suspension. ns: non-significant difference according to the Mann-Whitney-Wilcoxon two-sided test (p = 0.63). Mouse: N = 75; human: N = 127. **c.** Representative full-well images of mouse muscle fibers after 4 days of differentiation on fibrin substrates with different sizes of grooves, and on non-patterned controls. Magenta: myosin heavy chain. Scale bar: 1 mm. **d.** Distribution of mouse muscle fiber orientation on micro-grooved or control substrates. The different colors represent different replicates (n=3 wells per condition). **e.** Representative full-well images of human muscle fibers after 6 days of differentiation on fibrin substrates with different sizes of grooves, and on non-patterned controls. Magenta: myosin heavy chain. Scale bar: 1 mm. **f.** distribution of human muscle fiber orientation on micro-grooved or control substrates. The different colors represent different replicates (n=3 wells per condition). **g.** Proportion of muscle fibers aligned within ± 20° of the peak of the distribution (i.e. main orientation angle). Mouse: p = 7.99 ⋅ 10^-4^ ; human: p = 4.00 ⋅ 10^-4^. Mann-Whitney-Wilcoxon test. Grooved samples: N = 9; non-patterned samples: N = 6.

After 1-2 days in growth medium, the cells reached confluency and were transitioned to a differentiation medium, following previously optimized protocols.^39^ We changed the medium daily to promote fusion of myotubes and preserve cell viability and metabolic activity. After 4-6 days, thick fused muscle fibers were visible in all conditions. Tissues were fixed and stained with an antibody against myosin heavy chain typical of type II muscle fibers. A point-scanning confocal microscope was used to record and stitch together images of the full well for each condition (Figure 2 c-d), which were then used to quantify the distribution of muscle fiber orientations (Figure 2 e-f). Measuring the orientation at the scale of the whole well allowed us to account for potential biases that could arise from local alignment of muscle fibers. Irrespective of their origin, cells grown in grooved gels mainly aligned along the groove axis, while the orientations of cells grown on flat and unstamped gels were more variable (Figure 2 c-f). Since the groove size did not seem to affect the alignment efficiency, and the distributions of muscle fibers within flat and unstamped groups were also indistinguishable (Supplementary Figures 4 and 5), we pooled samples into two larger groups for subsequent analyses: grooved (12.5, 25, and 125 μm) versus non-patterned (flat and unstamped). To quantitatively verify that micro-grooved substrates enabled precisely controlling the alignment of muscle fibers, we measured the proportion of fibers whose orientation fell within ± 20° of the mean of the distribution of orientations (Figure 2g). For both mouse and human cells, the proportion of uniformly oriented cells was significantly higher in grooved samples than in the non-patterned ones (mouse: 49.9% ± 1.98% vs 32.9% ± 8.82%, p = 7.99 ⋅ 10-4 ; human: 28.7% ± 1.72% vs 19.5% ± 2.49%, p = 4.00 ⋅ 10-4). Moreover, a simple redesign of the stamp could be used to pattern aligned muscle in other culture formats, such as microfluidic devices (Supplementary Figure 6). Taken together, these results indicate that our one-step method enables simple templating of actuators via micro-topographical patterning (STAMP).

### STAMPed hydrogels influence muscle contraction direction, but not magnitude

Since the forces generated during muscle contraction are an important feature for functional characterization of engineered tissues for applications in high-throughput drug screening and robotics alike, we explored whether and how muscle function could be impacted by the micro-topography of the grooved hydrogel substrate. A function generator connected to a custom 24-well plate lid harboring electrodes was used to electrically stimulate mouse and human muscle layers grown on grooved and non-patterned fibrin following previously established methods.^42^ Brightfield microscopy videos of muscle contraction were recorded in response to electrical stimulation at 1 Hz (Supplementary Videos 1-10).

Using a custom open-source computer vision pipeline,^43–45^ we tracked tissue displacement with high 2D spatiotemporal resolution. We also calculated the displacement vectors between a contraction peak and the preceding relaxation phase in the presence or absence of grooves, enabling visualization of dynamic muscle contraction as vector fields (Figure 3a). Since muscle fibers generate force along their long axis, substrates with vertically aligned grooves yielded tissues that twitched in a more uniformly vertical direction than non-patterned substrates. As the total displacement that a contractile muscle layer generates is dependent on the direction and timing at which individual fibers contract, we calculated the mean absolute displacement (MAD) from each video and plotted its maximum (MADmax) for grooved versus non-patterned samples (Figure 3b). MADmax values were not statistically significant as determined by the Mann-Whitney-Wilcoxon two-sided test (mouse: p = 0.21; human: p = 0.39).

**Figure 3:**
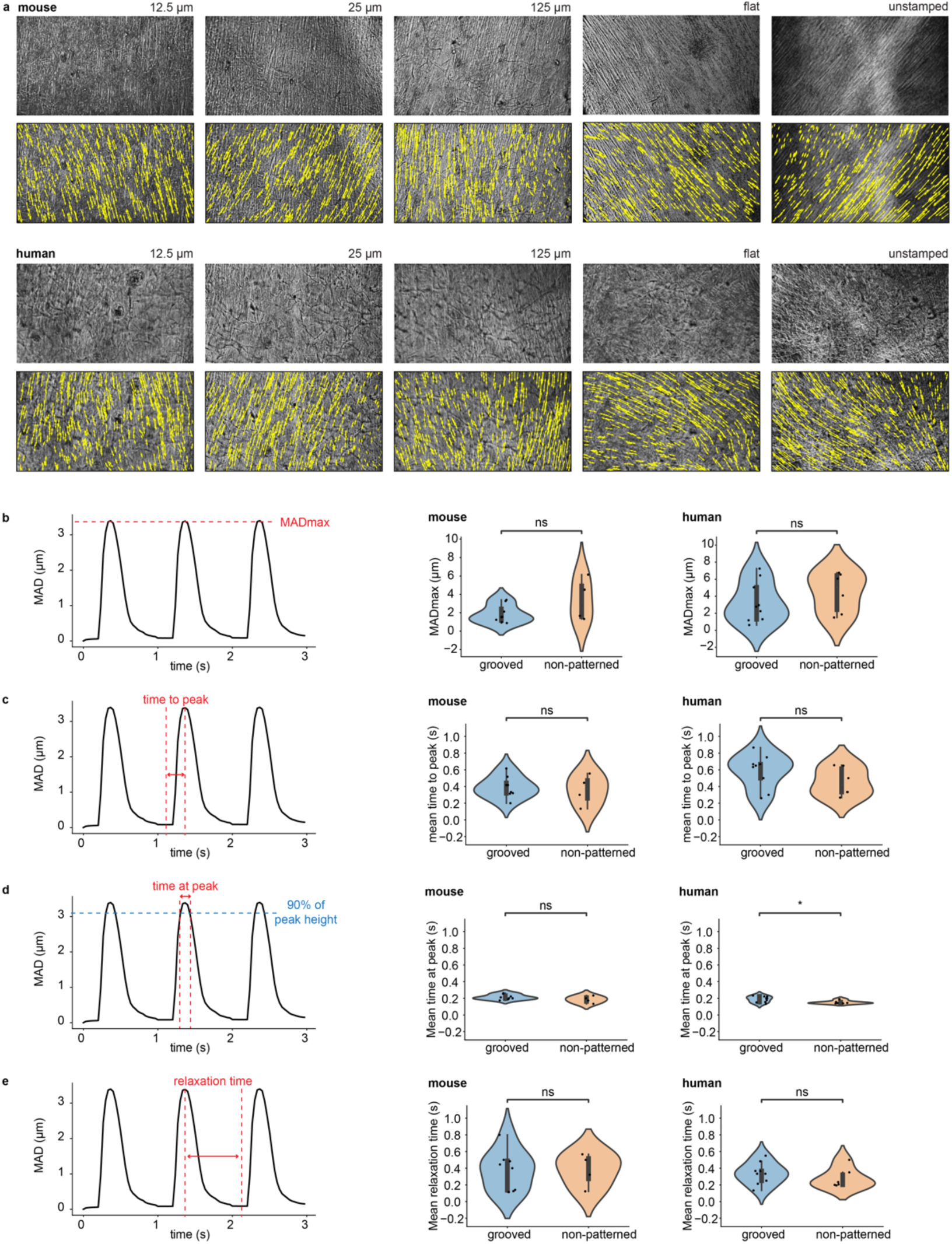
Impact of microscale topographical cues on muscle contraction. **a.** Representative spatial maps of displacement vectors generated by mouse and human cells upon electrical stimulation. Cells were either grown on substrates with microscopic grooves or non-patterned controls. Mouse and human cells were differentiated for 4 and 6 days, respectively. **b-e.** Functional metrics of muscle contraction achieved by mouse and human muscle fibers differentiated on grooved or non-patterned samples (mouse: day 4 of differentiation, human: day 6). These metrics include: **b.** the maximal value of the mean absolute displacement across the whole field of view (mouse: p = 0.21; human: p = 0.39). **c.** the time to peak, i.e. time it takes for the muscle fibers to contract from their relaxed state to the peak displacement value (mouse: p = 0.81; human: p = 0.26). **d.** the time at peak, defined as the time during which the mean absolute displacement was greater than 90% of the peak value (mouse: p = 0.37, human: p = 0.04955). **e.** the relaxation time from the peak back to the baseline MAD value (mouse: p = 0.86, human: p = 0.32). Throughout panels **b-e**, ns denotes a non-significant difference according to the Mann-Whitney-Wilcoxon two-sided test while * indicates p < 0.05. Number of biological replicates (wells) for: grooved mouse samples: N = 8; non-patterned mouse samples: N = 4. Grooved human samples: N = 9. Non-patterned human samples: N = 6.

We also explored the effect of grooved substrates on various metrics of contraction kinetics: time to peak force, time at peak force, and relaxation time, as these are critical to precisely control in the context of robotic actuators.^46^ The time to peak (Figure 3c), defined as the time it takes for a relaxed tissue to reach its maximal response to an electrical stimulus, did not vary significantly between grooved and non-patterned samples, irrespective of the species (mouse: p = 0.81; human: p = 0.26; Mann-Whitney-Wilcoxon test). Likewise, we analyzed the time at peak force, defined as the time during which the displacement was greater or equal to 90% of MADmax (Figure 3d). While for mouse tissues, there was no significant difference between the time at peak measured from grooved versus non-patterned samples (p = 0.37, Mann-Whitney-Wilcoxon test), the p-value obtained for human samples barely crossed the significance threshold of 0.05 (p = 0.04955). However, we note that such a small effect size is insufficient to warrant major interest, and could be a consequence of the relatively small sample size. Finally, we measured the tissues’ relaxation time (Figure 3e), the time during which the MAD decreased back to its relaxed state after a contraction peak. The presence of grooves did not significantly impact the relaxation time for either mouse (p = 0.86, Mann-Whitney-Wilcoxon test) or human samples (p = 0.32, Mann-Whitney-Wilcoxon test). Overall, these results indicate that the global fiber alignment imposed by grooves did not significantly impact muscle contraction kinetics.

Finally, we explored the effect of substrate topography on muscle maturation by quantifying the width of muscle fibers, their fusion index (a measure of how many nuclei are contained within each muscle fiber on average), and nuclei circularity (Figure 4). To this end, we stained myosin heavy chain and nuclei with MF20 and NucBlue, respectively, and acquired high-magnification confocal microscopy images. To calculate the fusion index (Figure 4a), we first segmented muscle fibers and nuclei before quantifying how many nuclei were contained inside a given muscle fiber. Fibers grown on grooved substrates had slightly higher fusion index values than those grown on non-patterned gels, but these differences were not statistically significant (mouse: p = 0.69; human: p = 0.08. Mann-Whitney-Wilcoxon test). We also quantified the width of muscle fibers by skeletonizing each segmented fiber (i.e. reducing it to a single line preserving the overall fiber orientation, as shown in Figure 4b) and measuring the shortest distance between each point of the skeleton and the edges of the fiber as detected by myosin heavy chain staining. The average distance multiplied by 2 for each fiber was defined as the fiber width. Fiber width differences were not significant between grooved and non-patterned samples irrespective of species (mouse: p = 0.14; human: p = 0.38. Mann-Whitney-Wilcoxon test). Of note, while mouse and human myoblasts had similar sizes upon trypsinization from their expansion flasks, the fibers they formed upon differentiation displayed different thicknesses (∼14 μm for mouse vs ∼9 μm for human). Finally, we also quantified the circularity of nuclei, as an additional metric to describe efficacy of myoblast fusion into multinucleated fibers. As anticipated, nuclei showed a reduced circularity compared to a perfect circle with a value of 1 irrespective of species and microtopography (no significant differences between grooved versus non-patterned samples - mouse: p = 0.11; human: p = 0.09. Mann-Whitney-Wilcoxon test).

**Figure 4:**
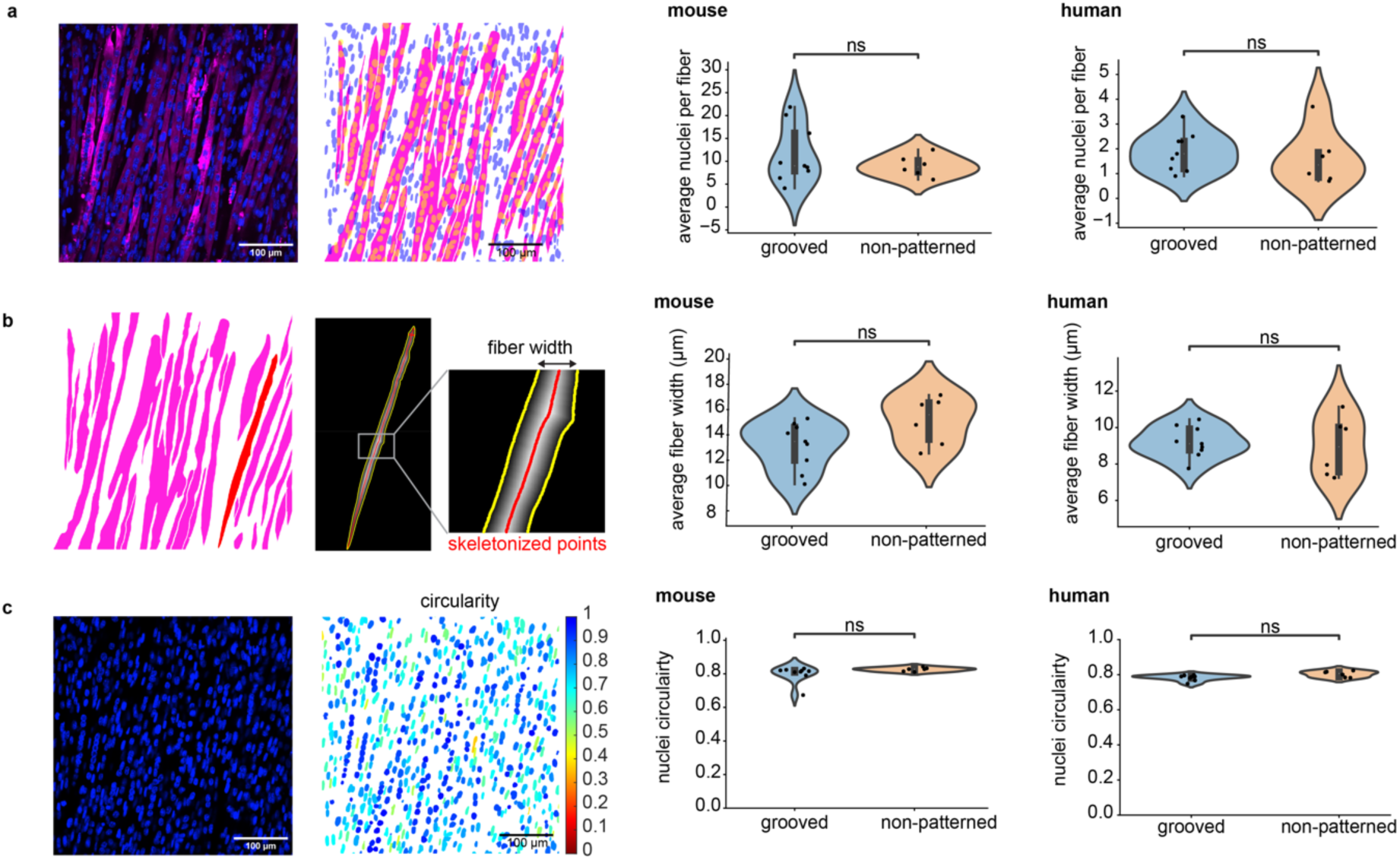
Impact of microscale topographical cues on muscle morphology and maturation. **a.** Representative images (stained and processed) and quantification of the number of nuclei per muscle fiber for mouse and human cells grown on grooved or non-patterned substrates (mouse: p = 0.69; human: p = 0.08). **b.** Width of mouse and human muscle fibers grown on grooved substrates or non-patterned controls (mouse: p = 0.14; human: p = 0.38). **c.** Nuclei circularity measurement (1: perfectly circular) for mouse and human cells on grooved or non-patterned substrates (mouse: p = 0.11; human: p = 0.09). Throughout the figure, ns denotes non-significant differences according to the Mann-Whitney-Wilcoxon two-sided test. Immunofluorescence pictures display myosin heavy chain in magenta and nuclei in blue, and 100 μm scale bars. Grooved samples: N = 9 biological replicates (wells). Non-patterned samples: N = 6 biological replicates (wells).

Taken together, these results indicate that tissue function and maturation is not affected negatively by the use of STAMPed hydrogel substrates, and that the direction and dynamics of engineered mouse and human muscle contraction can thus be precisely patterned with our one-step molding approach.

### Deploying STAMP to generate multi-oriented muscle actuators for planar soft robots

To demonstrate the versatility of STAMP, we showcased its potential for 2D spatial patterning by designing and manufacturing a planar soft robot featuring complex multi-oriented muscle architecture. Our robot was inspired by the architecture of the iris, in which a region of muscle circumferentially aligned around a pupil is surrounded by a region of radially aligned muscle.^47^ We computationally modeled our tissues as a bilayer composed of a thick fibrin hydrogel overlaid with a thin muscle monolayer, in which the muscle was represented by a thermal material with an expansion coefficient matching the strain generated by our engineered mouse muscle tissues on 25 μm grooves (∼6.7% at 1 Hz). This modeling setup enabled mimicking the spatially distributed forces generated by a muscle monolayer composed of thousands of individually contracting fibers.

Alongside the computational model, we designed a stamp of the same geometry, comprising a central region of concentric grooves surrounded by an outer region of radial grooves (Figure 5a). Similar to our multi-well plate patterning approach, we included a bubble release-feature to minimize structural defects arising from air trapped below the stamp. We then used the iris stamp to transfer the multi-oriented pattern onto a fibrin gel, as described and optimized above. STAMPed iris substrates were seeded with mouse myoblasts expressing the blue light-sensitive calcium ion channel channelrhodopsin2, conjugated to a TdTomato fluorescent tag.^16^ Upon differentiation, the cells formed muscle fibers that were precisely aligned to match the multi-oriented arrangement of the grooves, thereby mimicking the architecture of iris musculature (Figure 5b).

**Figure 5:**
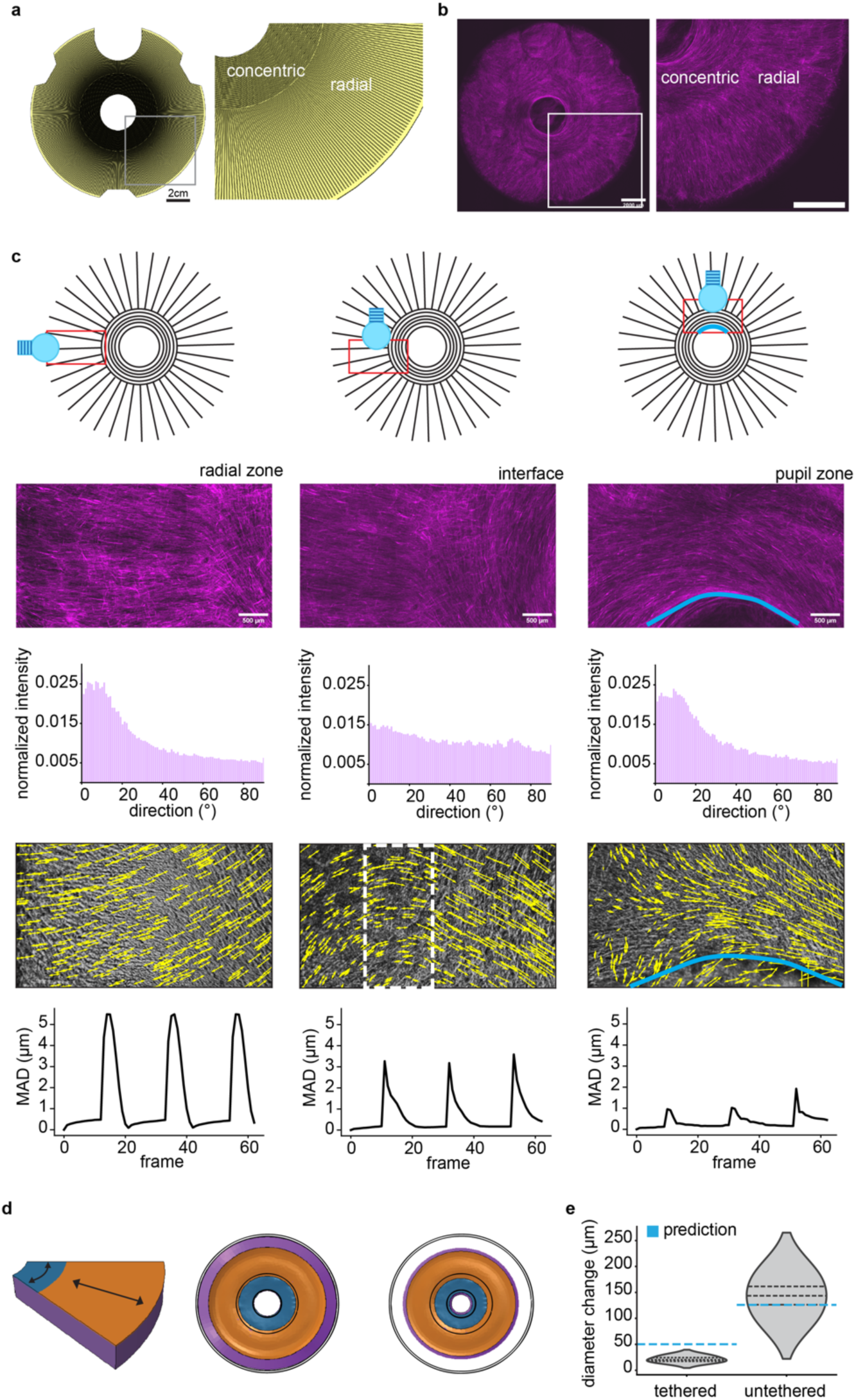
Leveraging hydrogel STAMPing to fabricate multi-oriented living actuators. **a.** CAD design of a micro-grooved stamp mimicking the multi-layered ring architecture of iris muscles. Scale bar: 2 cm. **b.** Mouse C2C12 cells growing on a fibrin substrate stamped with the iris-like pattern from panel a. Magenta: channelrhodopsin-TdTomato. Scale bar: 2 mm **c.** Schematic view of different regions of the iris, along with representative fluorescence images, directionality histograms, spatial maps of displacement vectors due to optically-stimulated muscle contraction at different locations of the well, and the corresponding mean absolute displacement signal (MAD). Regions of interest are symbolized by the red rectangle in the schematics, while the dashed white box indicates a transition zone where twitch direction shifts between the radial and concentric muscle layers. The outline of the pupil hole is shown in blue. The light bulb icon indicates the approximate position of the blue LED used for optogenetic stimulation. Scale bars: 500 μm. **d.** Left: 3D section of computational model of iris showing spatially segregated regions of concentric (blue) and radial (orange) aligned muscle monolayers on fibrin (purple). Model predicted displacement of the iris from its original configuration (denoted with black lines) is smaller in tethered (middle) than untethered configurations (right). **e)** Experimentally observed pupil diameter change in tethered and untethered irises as compared to computational model predictions.

Leveraging a custom fiber optic setup, we selectively stimulated different regions of the engineered iris with blue light (470 nm) to specifically trigger muscle contraction in only that region while recording brightfield microscopy videos (Supplementary Videos 11-13). Tracking data from our custom computational framework was used to plot spatial maps of the displacement generated by muscular contraction, as described above (Figure 5c). These analyses demonstrated that local displacement generally followed the direction of muscle fibers in that region, while highlighting transition zones at the interface between differently oriented regions. Moreover, we observed shrinkage of the pupil diameter in response to contraction of the multi-oriented actuator, mimicking the function of native iris musculature.

As biohybrid robots are often deployed in an “untethered” format to minimize resistance to contraction and maximize actuator stroke,^35,48^ we released the engineered iris from the underlying well plate and showcased a corresponding increase in the pupil shrinkage in response to light-triggered muscle contraction (Supplementary Video 14). Empirical results validated computational modeling of STAMPed irises in both tethered and untethered configurations (Figure 5d-e), though the model slightly overpredicted the pupil constriction of tethered irises. Overall, these results showcase the robustness of our method of designing, fabricating, and testing planar biohybrid robots capable of complex multi-DOF motion.

### Discussion and Conclusion

We have developed STAMP, a protocol for integrating microscale topography in extracellular matrix hydrogels, and demonstrated its ability to rapidly and precisely generate complex 2D patterns of aligned skeletal muscle tissues. Our method is compatible with a wide range of cross-linkable hydrogels as well as live cell microscopy, enabling high-throughput functional characterization of tissues via computational analysis of muscle contraction. In addition, it can be adapted to a variety of patterns beyond simple linear alignment, as demonstrated by the fabrication of an iris-mimicking multi-DOF bioactuator.

As compared to previous approaches of coating 3D micro-grooved scaffolds with extracellular matrix hydrogels, our method of generating aligned 2D muscle does not require complex multi-step manual handling or access to microfabrication facilities, and is also compatible with live cell imaging and untethered deployment in soft robots.^29–31^ Of note, there are other effective 2D muscle alignment strategies that are compatible with high-throughput imaging and longitudinal monitoring, such as growing myoblasts on electrospun nanofibers coated with adhesion-promoting biopolymers such as Matrigel.^5,20^ However, electrospinning requires custom experimental setups that are challenging to spatially control, thus providing limited precision over the 2D patterning of nanofibers. This limitation largely restricts electrospun substrates to fabricating unidirectionally aligned muscle layers which are suitable for high-throughput therapeutic screening (though not broadly accessible in all lab environments), but cannot be adapted to generate the multi-oriented architectures required for biohybrid robotics.

We have outlined a method to directly pattern complex microscale topography into natural hydrogel substrates with a one-step molding approach, enabling rapid patterning of complex multi-oriented muscle architectures. While others have also successfully patterned microgrooves into hydrogel scaffolds,^34,35,49^ these protocols involve long, multi-step fabrication processes, and rely on complex microfabrication tools that are not always readily available in resource-limited research institutions. In our study, we used a 3D printer to explore the effect of various groove sizes on muscle fiber orientation, and our analyses revealed that grooves as wide as 125 µm enabled reliable fabrication of aligned muscle tissues from both mouse and human cell sources.

This size is well within the print resolution of commonly accessible stereolithography 3D printers (SLA), and several companies even offer the printing and shipping of SLA-printed parts as a service. To facilitate open access and reproducibility of our method, stamp designs can also be virtually uploaded and readily shared between groups (see Supplementary Information for all our CAD files). Moreover, our simple optimized cleaning protocol facilitates mold reuse over multiple cycles, further increasing the cost- and time-effectiveness of STAMP. A downside of our one-step hydrogel casting and stamping process is that it increases the risk of trapping air bubbles in the polymerized hydrogel. Our bubble-release feature mitigates this problem, but microscale bubbles can still remain in the gel due to its viscosity. However, despite leading to minor local defects, we did not observe any impact on tissue global alignment due to the presence of micro-bubbles. Another drawback of our protocol is the dependence of pattern quality on uniform contact between the gel surface and the stamp grooves. This requires careful optimization of the gel volume and holder length when adapting STAMP to new cell culture formats, such as multi-well plates of different sizes or microfluidic chips (Supplementary Figure 6).

While we found that groove size did not significantly affect muscle fiber alignment in either our mouse or human cell-derived tissues, it is important to note that other researchers have identified “optimal” dimensions of microscale grooves^31,49^ or micropatterned ligands^26^ that can control the orientation of skeletal muscle tissue. These dimensions, however, vary across studies, ranging from less than 2 μm to 100 μm, suggesting that differences in the substrate fabrication process or in the quantification of alignment might influence the reported results.^30,49–52^ One explanation for this phenomenon is that most prior studies analyze alignment at a local scale (i.e. tissue sub-region spanning hundreds of microns), whereas we assess tissue-wide alignment across a region spanning several millimeters. Thus, our analyses may have disregarded subtle off-pattern angling sometimes reported for muscle fibers grown in wider grooves or ligand patterns.^26,49^ While these microscopic differences may be interesting to study, we consider global tissue alignment and coordinated contraction to be more useful metrics for both high-throughput drug screening and macroscopic biohybrid robots. Importantly, given our results with flat-stamped substrates, we do recognize there is a maximal groove width beyond which no muscle alignment would be observed. However, our results show that microgroove widths up to 125 µm are sufficient to coordinate the interplay of physical confinement and asymmetric cellular elongation required for efficient alignment of both mouse and human muscle. Future studies will be necessary to determine the roles of seeding density and growth rate, which must also influence the physical confinement of single cells.

Unlike groove width, the impact of groove depth on muscle orientation is rarely investigated. A recent study explored this interesting concept by fabricating silicon substrates for muscle with grooves of ‘touchable’ and ‘untouchable’ depths.^29^ They found that muscle fibers were unable to span gaps with untouchable depths (∼400 μm, i.e. around 20-30 times the size of single cells in suspension), which forced them to grow along the edges of the topological features. While we did not systematically probe the effect of groove depth in this study, we always observed confluent colonization of fibrin substrates, irrespective of microtopography. In other words, no groove depth we investigated was ‘untouchable’ for our cells. This difference could arise from material properties: as fibrin swells upon liquid absorption, the designed features lose fidelity, resulting in smoother edges, which may facilitate more uniform cellular colonization compared to the vertical walls of an etched silicon wafer. However, since we used stamps with equal width and depths to maintain equal aspect ratios in all conditions, additional investigations will be needed to determine whether changing the grooves’ aspect ratio influences muscle fiber orientation.

In our study, STAMP proved effective to align myoblasts from both human and mouse origin. However, others have shown that mouse and human myoblasts responded differently to substrates displaying micropatterned ligands.^26^ In their setup, C2C12 cells required a specific line spacing to align properly (20-30 μm), unlike primary human myoblasts, which they imputed to inter-species differences, such as the higher growth rate and inherent orientation bias they observed for C2C12 cells. In our study, the distributions of fiber orientation in grooved substrates also displayed different shapes between human and mouse cells, with better-defined peaks and narrower tails for mouse alignment distributions (Figure 2c-d). However, we attribute this difference to the noisier myosin staining for human cells, which impacted subsequent directionality analysis, rather than to inter-species variability. The presence of noise in the immunofluorescence pictures also lowered the proportion of fibers oriented within ±20° of the mean by introducing randomly oriented speckles. Even so, the alignment of muscle fibers was significantly more uniform when they grew on grooved-versus non-patterned fibrin, irrespective of the species of cellular origin (Figure 2e).

Our results also demonstrate that the presence of grooves did not significantly impact muscle differentiation or strength from either mouse or human cells, as assessed by both morphological metrics (muscle fiber width, fusion index, nuclei circularity) and functional metrics (maximum displacement, contractile dynamics). By contrast, other studies have found that alignment promotes muscle fiber differentiation. For example, studies of C2C12 cells grown on grooved elastomeric thin films have shown an increase in fusion index and expression of differentiation markers as compared to flat thin films.^31^ However, it is important to note that these studies focus on regions of local alignment on short length scales (hundreds of microns), and we and others have noted that skeletal muscle grown on compliant substrates will exert traction forces to align locally even in the absence of grooves,^43^ likely driving the observed increase in differentiation markers. By contrast, skeletal muscle grown on flat stiff substrates such as glass demonstrate much less local alignment (Supplementary Figure 7). Thus, we hypothesize that *local* alignment of fibers (which happens spontaneously on soft substrates) is required to promote muscle differentiation and *global* alignment of muscle fibers (which requires micropatterned cues) is required to control macroscale motion. STAMP enables controlling both local and global alignment with high fidelity.

In the future, STAMP could be readily adapted to a range of cell culture formats, such as wells with different diameters, Petri dishes, or even microfluidic chips, simply by editing the design of the 3D printed parts (Supplementary Figure 1). Because STAMP is independent of hydrogel chemistry, it could also be used to pattern a variety of ECM-mimicking hydrogels suitable for different cell types such as collagen or gelatin methacrylate, as long as the crosslinking kinetics enable casting and stamping of the liquid prepolymer prior to sol-gel transition. Our demonstration of using STAMP to align cells of both mouse and human origin adds to published research showing that micro-grooved substrates enable controlling the orientation of various cell types. Beyond skeletal muscle and neuromuscular junctions,^50,53^ microgrooves have also been shown to enhance the alignment of cardiomyocytes,^54^ neurons,^55,56^ and periodontal ligament cells,^57^ among others. However, as STAMP does not enable precise positioning of single cells upon seeding, adaptation of the method towards cell types that require specific initial configurations (such as engineered neuron networks) will require combining with other existing methods for spatially controlling cell seeding.^58,59^

Finally, STAMP’s ability to reproducibly generate high-fidelity high-resolution muscle fiber patterning enabled us to design a planar soft robot capable of complex multi-DOF motion in response to muscle contraction. In previous demonstrations of biohybrid robots actuated by 3D skeletal muscle, the engineered bioactuator contained largely unidirectionally aligned fibers and was thus only capable of 1-DOF contraction. While these actuators have been sufficient to demonstrate proof-of-concept applications such as walking or pumping or swimming,^9^ there remains a need to design and deploy engineered muscle actuators that reproduce more complex architectures and motion patterns as observed in native tissues. Likewise, while planar configurations for biohybrid robots have proven very impactful for machines powered by 2D sheets of cardiac muscle,^21–25,32,60^ there have been fewer demonstrations of skeletal muscle monolayers cultured on untethered 2D substrates,^61,62^ likely due to the large passive and active tensions generated by individual fibers that lead to cell delamination and wrinkling of substrates made from polymer thin films. In this study, we leverage fibrin as a substrate for a planar soft robot, as fibrin enables longitudinal 2D culture of skeletal muscle over several weeks without delamination or wrinkling.^41^ We deploy STAMP to generate a complex micro-topographical pattern in fibrin that yields a multi-oriented muscle actuator capable of multi-DOF motion. These combined insights enabled recreating the action of an iris containing regions of concentric and radially aligned muscle capable of regionally-specific force generation. Moreover, leveraging optogenetic muscle cells enabled us to generate complex behaviors such as light-controlled pupil constriction, as validated by computational modeling. We anticipate that the accessibility and reproducibility of our STAMP technique, combined with our method for predictive computational modeling of planar soft robots actuated by skeletal muscle monolayers, will enable others in the field to design and deploy 2D biohybrid robots capable of a wide range of programmable multi-DOF actuation behaviors.

## Methods

### Stamp manufacturing

All the stamp holders described in this work were designed using a CAD software (Solidworks) and exported as .stl files to use with Formlab’s Preform Slicer software. The stamp holders were printed out of Surgical Guide V1 Resin without supports and with a layer height of 50 μm using a Formlabs 3B Stereolithography (SLA) printer (Formlabs). The stamps were post-processed in a Form Wash with Isopropyl Alcohol for 45 minutes and cured in a Form Cure UV curing chamber for 30 minutes at 70 °C as recommended by Formlab’s Surgical Resin user guide^63^. The stamp holders were further cured at 80 °C in an oven for at least 20 hours to cure any remaining resin before combining them with UpPhoto micro-grooved stamps (described below). Just before attaching the micro-grooved stamps, the stamp holders were washed with soapy water, dried, and then run through a Tuttnauer benchtop line autoclave (Tuttnauer USA Co Ltd, 2840EL-D) at 121°C using the preset plastic cycle and allowed to cool to room temperature.

Micro-grooved stamps (shown in Figure 1a-c, Figure 5a, and Supplementary Figure 4) were designed using Solidworks and exported as .stl files to use with UpNano’s slicer software Think3D (<u>UpNano GmbH, Vienna, Austria</u>). The flat stamps and the stamps with 125 μm feature sizes were printed using a 5x objective lens and UpPhoto Resin on an UpNano One 2-photon printer (UpNano GmbH). Stamps with smaller feature sizes of 12.5 μm and 25 μm including the 25 μm ocular stamps, were printed with UpPhoto Resin on a 10x objective (<u>UpNano GmbH</u>). All stamp prints were post-processed by progressively soaking each print in 3 different isopropyl alcohol baths for 10 minutes each and then removing the prints from the build plate using a razor. All stamps were allowed to sit and further photo cure for at least 12 hours to remove any remaining uncured resin before attaching the UpPhoto micro-grooved stamps to the Surgical V1 stamp holders described above. The UpPhoto stamps were glued to the stamp holders using KwikSil™ silicon adhesive (World Precision Instruments). Each 24-well plate stamp was aligned using the port hole as a clocking feature in addition to kinematic pins and notches included in the design that assist in centering the stamps on the stamp holder. For all stamps, the silicon adhesive was allowed to set for 7 minutes before sterilization. Stamps were sterilized under UV light in a biosafety cabinet for 30 minutes before each use. In addition, between each use, stamps were cleaned as described in Stamp Reuse.

### Stamp use and gel preparation

To facilitate subsequent release, the stamps were first soaked for 1 hour in bovine serum albumin (BSA, Thermo Fisher Scientific, 37525) diluted to 1% concentration. The alignment stamps and ocular stamps were placed in 24-well plates (Avantor, 10891-558), each with 1.5 mL of BSA. After 1 hour, the stamps are removed from the BSA bath and allowed to dry to remove excess BSA. Once dry, the stamps are placed in ultrapure milli-Q water (Sigma Aldrich, ZRQSVR3WW) just before using them to micropattern and stamp fibrin gels in 24-well plates. The fibrin gels were made at the same concentration as described in Bu et al.^41^. We created a solution of 8 mg of fibrinogen from fetal bovine plasma (Sigma Aldrich, F8630-1G) and 0.4 uL of thrombin from bovine plasma (Sigma-Aldrich, T4648-1KU) per mL of Dubelco’s Modified Eagle Medium (DMEM) (Sigma Aldrich, D6429-500ML). 590 μL of the mixture of fibrinogen and thrombin was then pipetted into each well of a 24-well plate (Cellvis, P241.5HN) before stamps were added. The stamps were then placed into the wells, in contact with the gel, and the plate was lightly tapped and tilted to push any visible air bubbles trapped underneath the stamp towards the bubble-release feature. After loading and stamping, the platforms were all placed in an incubator at 37 °C for 1 hour. After incubation and polymerization, the stamps were carefully removed and the gels were kept hydrated with 0.5 mL of growth medium until they were seeded with cells.

### Stamp reuse

After the stamps described in section 1 were used to pattern gels, they were cleaned by first rinsing and soaking them in water immediately after use. They were then sonicated for at least 20 minutes in an isopropyl alcohol bath and dried using lens cleaner paper to remove any gel residue from the microgrooves. The stamps were then sterilized under UV light in a biosafety cabinet for 30 minutes before reuse. Pictures of gels made with stamped used for the 4th time after being cleaned can be found in Supplementary Figure 3. We did not observe any significant difference between gels made with stamps used for the first time and those used for a fourth time after cleaning.

### Cell culture

Mouse myoblasts C2C12 cells were used both for linear alignment studies (C2C12 WT cells, ECACC, 91031101-1VL) and for the multi-oriented iris-mimicking construct (optogenetic C2C12 cells expressing 470 blue light sensitive Channelrhodopsin [ChR2(H134R)] tagged with tdTomato ^16^. They were expanded in growth medium (GM) consisting of fetal bovine serum (Life Technologies, A5670701), Corning™ L-glutamine solution (Fisher Scientific Co LLC, MT25005CI), and 1% penicillin-streptomycin (Fisher Scientific Co LLC, MT30002CI) dissolved in DMEM at ratios of 1:10 vol/vol, 1:100 vol/vol, and 1:100 vol/vol respectively. The cells were seeded at a density of 100,000 cells per well of a 24-well plate and grown in growth medium (GM) with 6-Aminocaproic acid (ACA) (Sigma-Aldrich, A2504-100G) at a ratio of 1:50 vol/vol for one day until they reached confluency. They were then switched to differentiation medium (DM) consisting of horse serum (Gibco, 26050-088), Corning™ L-glutamine solution (Fisher Scientific Co LLC, MT25005CI), and 1% penicillin-streptomycin (Fisher Scientific Co LLC, MT30002CI) dissolved in DMEM at ratios of 1:10 vol/vol, 1:100 vol/vol, and 1:100 vol/vol respectively with ACA and inuslin-like growth factor-1 (IGF-1) (PeproTech, 100-11R3-1MG) added to the whole solution at a ratio of 1:50 vol/vol and 1:20000 vol/vol respectively. The cells were allowed to differentiate with media replacement every other day before being electrically stimulated, fixed, and stained, as described below.

Passage 2 Human skeletal muscle myoblasts (Cook Myosite, #SK-1111-P01547-29M) were seeded on hydrogels at a density of 100,000 cells per 24-well plate. The skMDC cells were grown in growth media consisting of MyoTonic Basal Medium (Cook Myosite, MB-2222) and MyoTonic Growth Supplement (Cook Myosite, MB-3333) supplemented with 1% penicillin-streptomycin. After 1 day the cells were switched to differentiation media consisting of KSR/CHIR/TGFbi/prednisolone [KCTiP] as described elsewhere^64^ supplemented with 0.2% penicillin-streptomycin and 2% ACA.

On the 6th day after switching to differentiation media, the cells were electrically stimulated, fixed and stained as described below.

### Electrical stimulation

Electrical stimulation was carried out by placing the cultures in 2 mL of DMEM and then supplying a 1 Hz 10 Vpp 20% duty cycle square wave using a Siglent SDG1032X function generator connected to electrodes made of platinum wires (Thermo Fisher Scientific, 010286.BY) suspended approximately 7 mm apart and 10 mm above the bottom of the well plate. The cultures were each stimulated for 10s and their twitch response was monitored with a brightfield Zeiss Primovert microscope equipped with an Axiocam 202 mono camera (Zeiss). Videos of muscle contraction were recorded at 30 frames per second at the center of each well in between the two electrodes, and 3 seconds were extracted from each video to quantify displacement magnitude, kinetics and direction, as described below.

### Optical stimulation

The optogenetic C2C12 cells used for the iris cultures were stimulated with both the same electrical setup described above and optically using 1 Hz blue light stimulation with a 20% duty cycle through a 470 nm optical fiber (Thorlabs), powered by an Agilent E3630A power supply (Agilent Technologies). Videos of muscle contraction in response to optical stimulation were acquired at 30 frames per second with a Zeiss Primovert Microscope.

### Video tracking and analysis

Spatial maps of displacement were generated using a freely available tracking algorithm as described in previous studies ^43–45^. Among other features, this code returns the X and Y positions of the automatically-identified tracking features over the whole duration of the video, along with their mean absolute displacement (MAD) in each frame. From these results, we calculated and plotted various functional metrics of tissue contraction in Jupyter Notebooks ^65^ using the following packages: NumPy ^66^, SciPy ^67^, Pandas ^68^, Matplotlib ^69^, Seaborn ^70^, cv2 ^71^. MADmax was quantified by calculating the maximal value of the MAD time series for each 3-second video of electrically stimulated, linearly aligned muscle fibers. Time to peak and relaxation time were obtained by identifying the peaks and valleys of the MAD signal (using ‘scipy.signal.find_peaks’) and calculating the time difference between a valley and the following peak, or between a peak and the following valley. We defined the time at peak as the time during which the MAD value was ≥ 90% of the peak value.

To generate spatial maps of displacement vectors, we used the X and Y positions of tracking features extracted from video frames corresponding to one valley of the MAD signal and the peak immediately following it. Because of the large number of tracking features, we downsampled each position vector by a factor 25. We calculated the X and Y displacement vectors by subtracting the X and Y positions of the initial time frame from the corresponding positions in the final time frame. We then used a combination of matplotlib.pyplot’s function ‘quiver’ and cv2’s ‘imshow’ to display the displacement vectors overlaid with a brightfield image of the muscle fibers that generated this displacement. Note that, to emphasize the bi-directional nature of muscular contraction and relaxation, we plotted each vector in both its positive and negative direction, which resulted in double-sided arrows.

### Immunohistochemical staining and fluorescence imaging

After stimulation, electrical or optical, cells were fixed by soaking cultures in 4% paraformaldehyde solution (Avantor, 100503-917) dissolved in phosphate-buffered saline (PBS) (Thermo Fischer Scientific, 20012027) for 15 minutes. The paraformaldehyde was then washed away with three rinses of PBS. Tissues were then permeabilized using 0.5% Triton X-100 (Sigma Aldritch, T8787-50ML) for 20 minutes followed by three 5-minute PBS washes. Cultures were then blocked with 1% bovine serum albumin in PBS for 45 minutes. The muscle tissue cultures were then stained with 150 uL 1:100 Myosin4 antibody conjugated to eFluor™ 660 (Invitrogen, 50-6503-82) for 1 h. Cultures were then washed with PBS three times and stained with NucBlue (Thermo Fischer Scientific, R37605) for 15 minutes following the manufacturer’s instructions. After one final round of three PBS washes, 3 drops of antifade media were added to each well. The C2C12 Channelrhodopsin2-expressing cultures constitutively expressed a TdTomato tag and were therefore not stained.

Following staining, all cultures were imaged using a Nikon AXR point scanning confocal microscope. To create the full-well images shown in Figures 2 and 4, we stitched together maximum intensity projection images obtained from z-stack images acquired across the entire well using a 4X objective. To acquire higher magnification pictures, which involve objectives with lower working distances, we carefully extracted the fibrin gels (along with the fixed cells on top) from the 24 well plates using a spatula. We then flipped them over and transferred them to glass-bottom 6-well plates (Cellvis, P06-14-1.5-N) and imaged them with a 40x objective.

### Image processing

Prior to calculating directionality spectra, the full well stitched images were binarized and filtered using MATLAB’s imbinarize() function with an adaptive filter and by removing any particles in the image with an area less than 1000 px^2^ for the mouse tissue culture images and 100 px^2^ for the human tissue culture images with bwareafilt() from MATLAB’s image processing toolbox ^72^. Directionality distributions were then calculated by applying the Directionality function from ImageJ Fiji^73^ to a centered 8 mm by 12 mm cropped region of the filtered stitched full well images. This region of interest captured the majority of the well, with the exception of the bubble trap and kinematic features at the edges of the well. Bin sizes of 2.022 degrees were used and the percent of fibers within ±20° of the mean direction for each replicate was computed by summing the normalized values of the polar Fast Fourier Transform power-spectral density provided by the ImageJ Directionality plug-in^73^. The distributions shown in Figure 1b and 1d were plotted using an adjusted version of the superviolin code package described in Lord et al. 2020. ^74^

Immunofluorescence images acquired at 40X magnification from the center of each well were used to quantify the fiber width and the number of nuclei per fiber. The C2C12 mouse fiber fluorescent channels were segmented into individual fibers by hand tracing the edges of the stained fibers in each image using Procreate drawing software^75^ on an iPad with an Apple Pencil, and human fibers, since the stains used in this work did not resolve the human fibers as well as C2C12 fibers, were segmented based on pseudo-brightfield images provided by the confocal. The nuclei fluorescent channels stained with NucBlue for both cell types were segmented by using the Cellpose nuclei identification package^76^. A nucleus was considered inside a muscle fiber if at least 50% of its segmented area overlapped with that of a given muscle fiber’s area (see figure 4a). The fiber width for each fiber segment within the field of view was calculated by skeletonizing each fiber and calculating the shortest distance to the edge of the segmented fiber from each skeleton point making use of the MATLAB functions bwskel() and bwdist()^72^. The final reported width for each segmented fiber was then calculated by multiplying all the measured distances to the skeleton points by 2 and then taking the average value for each fiber. The average nuclei circularity of each biological replicate was reported using the MATLAB function regionprops() to calculate a circularity value between 0-1 for each segmented nuclei in each replicate^72^. The average circularity for each replicate is reported in figure 4c.

### Statistical analysis

Directionality, contraction, and morphological metrics from grooved and non-patterned samples were compared using the Mann-Whitney-Wilcoxon statistical test with the Statannotation Python package. ^77^ P-values < 0.05 were considered statistically significant.

### Computational modeling

To model the contraction of the grooved fibrin/ iris muscle, a thermal load simulation was created to mimic the strain applied by the muscle during contraction. First, the entire bilayer was created using Abaqus finite element analysis software. From there, the part was partitioned twice: once to separate the muscle layer from the fibrin (creating a 50um micron thickness for the muscle layer and a 1mm thickness for the fibrin) and twice to partition the radial grooved section from the circumferential section. In the property module, three elastic materials were created all with ∼0.5 Poisson’s ratio. The fibrin was entered as an elastic material with Young’s module of 0.3kPa and the two muscle materials were entered with a Young’s modulus of 130kPa, based on our previously published evaluations of fibrin and muscle mechanical properties.^41,46^ Two muscle materials were created to assign different thermal expansion directions to mimic the directionality of the stamped grooves. The circumferential muscle material was assigned a 0.067 thermal expansion value parallel to the force-generating axes of the fibers, and the same value was assigned to the radial muscle material in their corresponding fiber direction. The coordinate system was changed to cylindrical for the creation of these orthotropic materials. The 0.067 alpha value corresponds to 6.7% strain for a 1 °C temperature drop. This strain value was determined experimentally from video analysis by taking the normal and shear strains over the iris during contraction at 1Hz, 10v, 20% duty cycle, and calculating the maximum principal strain. An axisymmetric constraint about the z axis, as well as a zero-displacement constraint in the z axis for the bottom surface of the fibrin layer, were used for both the tethered and untethered models. For the tethered model, x and y displacement constraints were also added. 1900 hybrid linear 3D elements were used to mesh the entire iris, and a 1 °C temperature drop was imposed to invoke a contractile strain response in the muscle layer.

## Author contributions

Conceptualization: T.R., L.S., R.R.; Data curation: T.R., L.S.; Formal analysis: T.R., L.S., M.B.; Funding acquisition: R.R.; Investigation: T.R., L.S., P.U.; Methodology: T.R., L.S., M.B., R.R.; Project administration: R.R.; Resources: R.R.; Software: T.R., L.S., M.B.; Supervision: R.R.; Validation: T.R., L.S., M.B. R.R.; Visualization: T.R., L.S., M.B., R.R.; Writing – original draft: T.R., L.S., M.B. R.R.; Writing – reviewing and editing: T.R., L.S., M.B. R.R.

## Data availability

The data supporting this article have been included as part of the Supplementary Information.

## Acknowledgements

The authors would like to thank Ayelet Lesman and Roi Habba for technical discussions regarding computational modelling. This work was supported in part by the US DoD Army Research Office Early Career Program (awarded to R.R), NSF CAREER Award (awarded to R.R.), US DoD Office of Naval Research Young Investigator Program (awarded to R.R.), the US DoD DURIP Program (awarded to R.R.), the La Caixa Postdoctoral Fellowship at MIT (awarded to T.R.), and the NSF Graduate Research Fellowship Program (awarded to M.B.). This this work was carried out in part through use of MIT.Nano’s facilities.

