## Supplementary Information for "Leveraging microtopography to pattern multi-oriented muscle actuators"

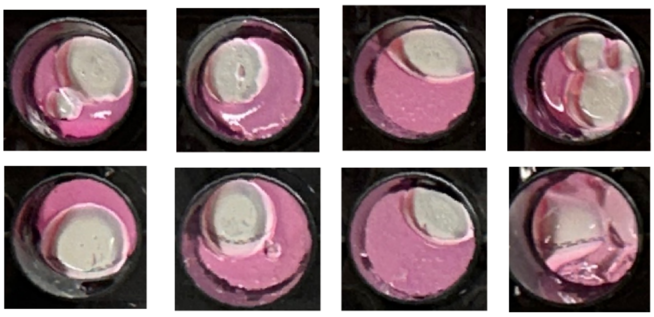

**Supplementary Figure 1:** Images of STAMPed fibrin gels in wells where the stamp did not have a “bubble release” feature. The lack of a bubble release feature resulted in large voids in the gels regardless of the amount of gel added to the well.

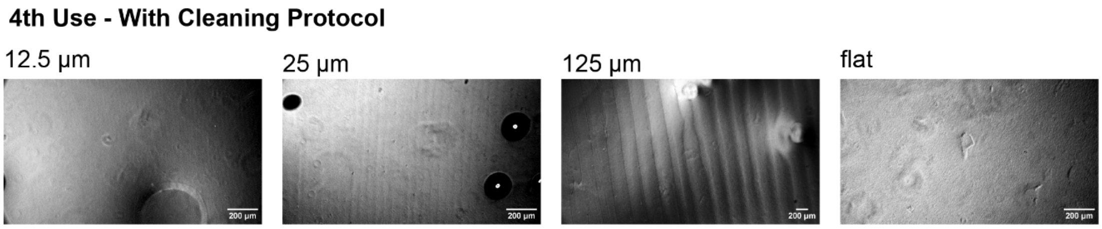


**Supplementary Figure 2:** Brightfield images of fibrin gels fabricated with stamps that were sonicated and cleaned as described in the methods section after being used 3 times previously. We observed no major differences in the visual quality of the grooves made by the reused stamps.

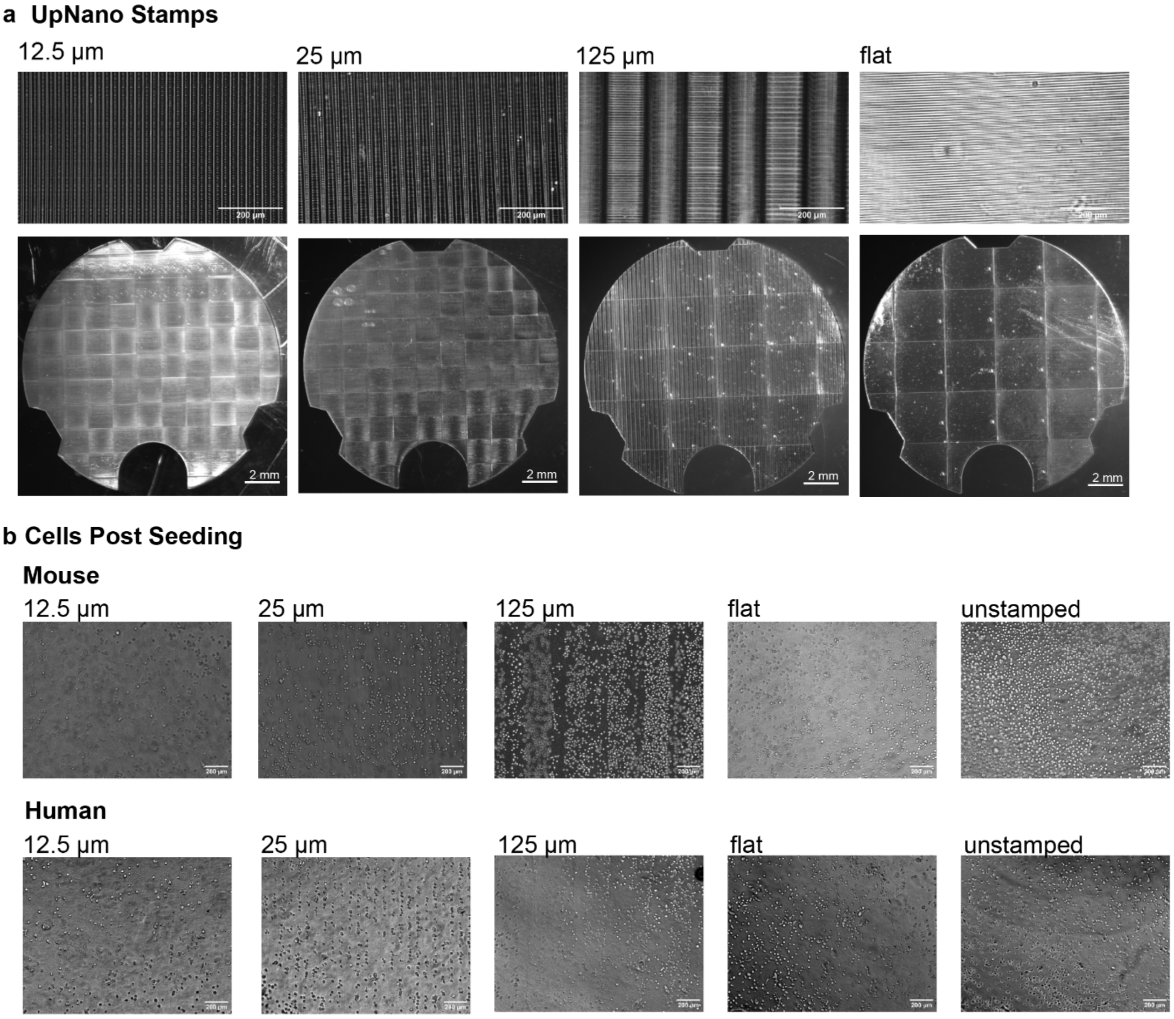


**Supplementary Figure 3. a)** 20x magnification brightfield images of each type of stamp feature size used in this study and corresponding stereoscope images of the entire stamp. The small parallel lines seen in the 20x brightfield images reveal the resolution of the UpNano One printer and the large uniform square lines in the stereoscope images show the working field of view of the different UpNano One objectives, along with the regions of overlap used to stitch the full stamp together during printing. The stamps with 12.5 μm and 25 μm grooves stamps were printed with a 10x objective and the 125 μm and flat stamps were printed with a 5x objective, resulting in a larger field of view and wider stitching regions. **b)** Brightfield images taken with a 10x objective showing initial cell seeding patterns under all the different gel patterning conditions used in this study.

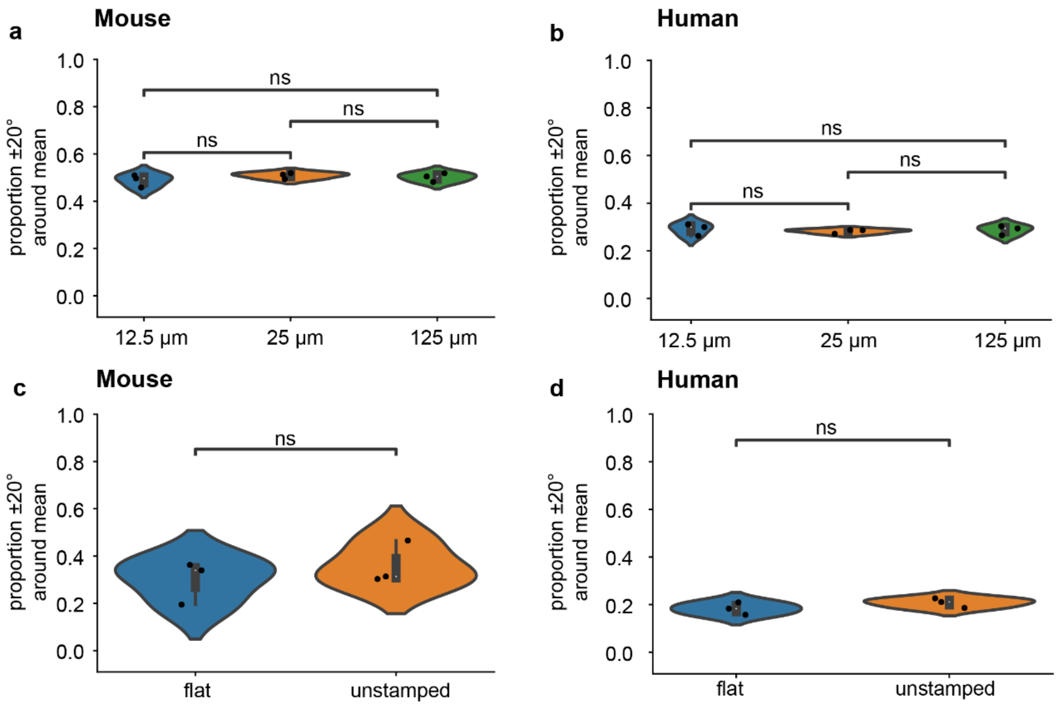


**Supplementary Figure 4:** Comparisons of the proportion of fibers within 20° of the mean of the fiber orientation distribution calculated as described in the main text. The source data are identical as in Figure 2g, which presents them pooled in two broader categories: grooved and non-patterned. **a)** Comparison of the proportion of C2C12 fibers within 20° of the mean direction between the different grooved feature sizes. We found no significant difference between groups with a Kruskal-Wallis test ($p = 0.49$). **b)** Comparison of the proportion of human fibers within 20° of the mean direction between the different grooved feature sizes. We found no significant difference between groups with a Kruskal-Wallis test ($p = 0.73$). **c)** Comparison of the proportion of C2C12 fibers within 20° of the mean direction between the different non-patterned substrates. We found no significant difference between the flat and unstamped control groups with a Kruskal-Wallis test ($p = 0.83$). **d)** Comparison of the proportion of human fibers within 20° of the mean direction between the different non-patterned substrates. We found no significant difference between the flat and unstamped control groups with a Kruskal-Wallis test ($p = 0.13$).

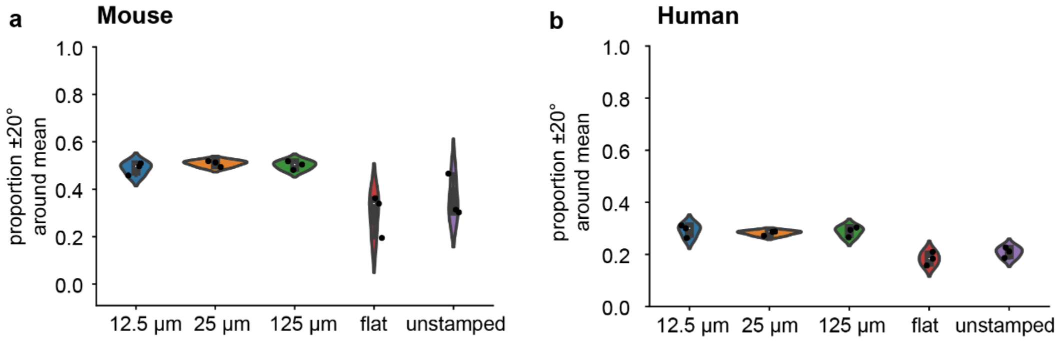


**Supplementary Figure 5:** The same data from Supplementary Figure 5 displaying all stamp sub-conditions on the same axis, reflecting the results from figure 2g in the main text. **a)** Proportion of C2C12 fibers oriented within 20° of the mean orientation across all gel feature size groups in both the grooved and non-patterned conditions. **b)** Proportion of human fibers oriented within 20° of the mean orientation across all gel feature size groups in both the grooved and non-patterned conditions.

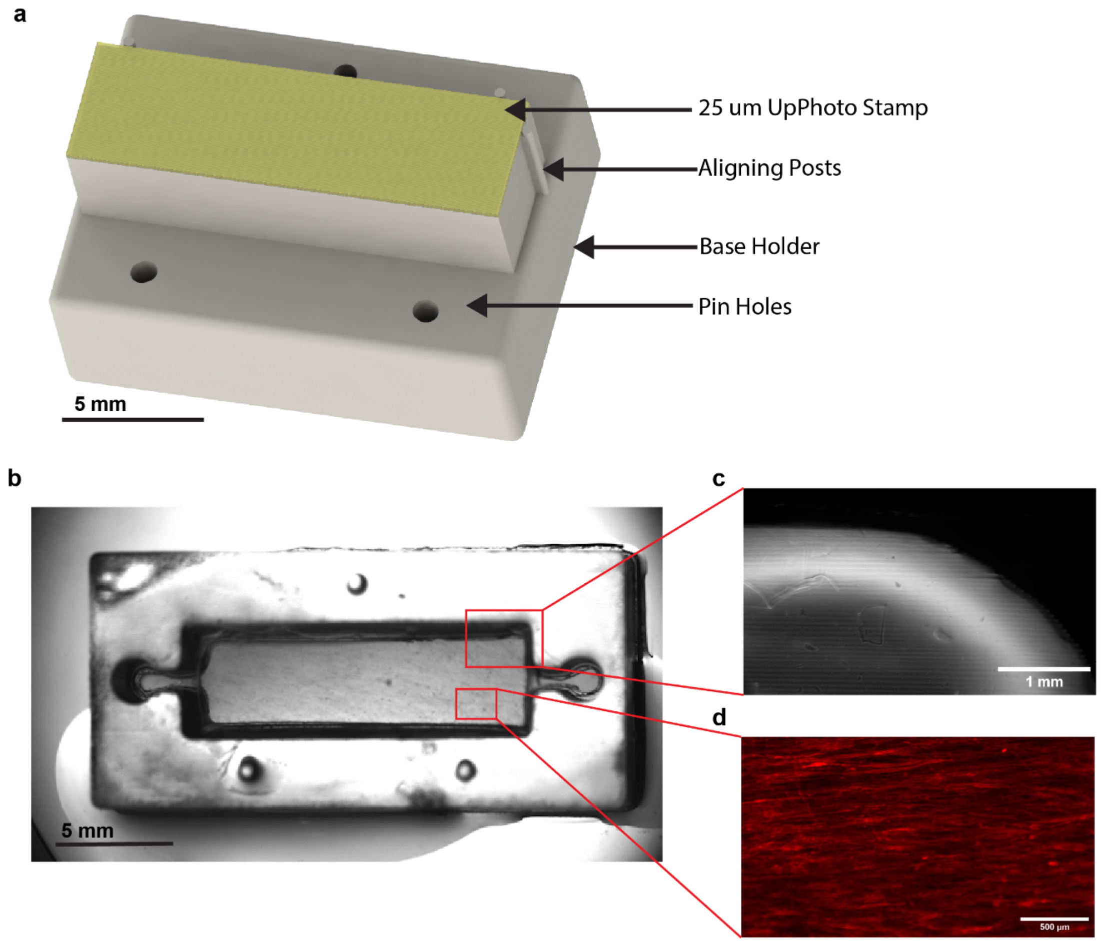


**Supplementary Figure 6: a)** CAD rendering of the stamp used to pattern the gel in the microfluidic chip shown in (b). A resin base aligns to the pins on the chip in (b) and holds an UpPhoto stamp adhered with KwikSil™ and aligned to the base with 3 aligning posts **b)** brightfield stereoscope image of a resin microfluidic chip to grow skeletal muscle under perfusion of culture medium. **c)** brightfield image of patterned gel right before cell seeding in the chip. The chip gel was patterned using a rectangular stamp with 25 µm grooves. **d)** fluorescence micrograph of optogenetic C2C12 cells grown in the stamped chip on their 6th day after being placed on differentiation media. Red: tdTomato-tagged channelrhodopsin.

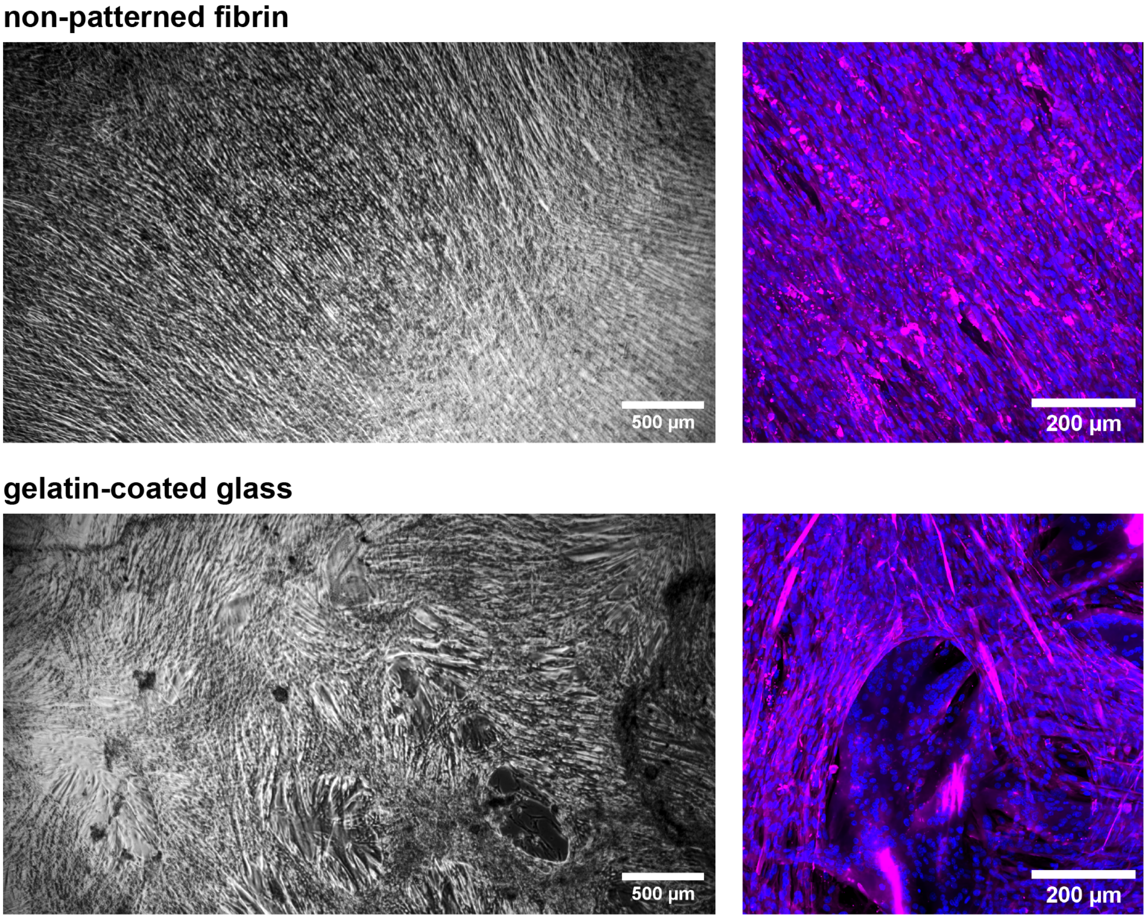


**Supplementary Figure 7:** Brightfield and confocal microscopy fluorescence images of mouse skeletal muscle grown on non-patterned fibrin gel and on glass coated with a 2% gelatin solution. Cells were imaged in brightfield after 3 days of differentiation, and fixed after 5 days of differentiation, prior to immunohistochemical staining and confocal imaging. Magenta: MYH2 stained with mouse 1F1B6 antibody. Blue: nuclei stained with NucBlue.

**Supplementary Videos and CAD Files can be found at this link.**

**List and description of supplementary videos:**

- **Supplementary Video 1:** electrically-stimulated contraction of murine myofibers grown on fibrin with 12.5 µm microgrooves.
- **Supplementary Video 2:** electrically-stimulated contraction of murine myofibers grown on fibrin with 25 µm microgrooves.
- **Supplementary Video 3:** electrically-stimulated contraction of murine myofibers grown on fibrin with 125 µm microgrooves.
- **Supplementary Video 4:** electrically-stimulated contraction of murine myofibers grown on a non-patterned fibrin gel molded in contact with a flat stamp.
- **Supplementary Video 5:** electrically-stimulated contraction of murine myofibers grown on non-patterned, unstamped fibrin gel.
- **Supplementary Video 6:** electrically-stimulated contraction of human myofibers grown on fibrin with 12.5 µm microgrooves.
- **Supplementary Video 7:** electrically-stimulated contraction of human myofibers grown on fibrin with 25 µm microgrooves.
- **Supplementary Video 8:** electrically-stimulated contraction of human myofibers grown on fibrin with 125 µm microgrooves.
- **Supplementary Video 9:** electrically-stimulated contraction of human myofibers grown on a non-patterned fibrin gel molded in contact with a flat stamp.
- **Supplementary Video 10:** electrically-stimulated contraction of human myofibers grown on non-patterned, unstamped fibrin gel.
- **Supplementary Video 11:** optically-stimulated contraction of optogenetic murine myofibers in the radially-aligned zone of an iris-mimicking multi-oriented muscle.
- **Supplementary Video 12:** optically-stimulated contraction of optogenetic murine myofibers at the interface between the radially and concentrically aligned zones of an iris-mimicking multi-oriented muscle.
- **Supplementary Video 13:** optically-stimulated contraction of optogenetic murine myofibers in the concentrically-aligned zone of an iris-mimicking multi-oriented muscle, at the edge of the pupil hole.
- **Supplementary Video 14:** shrinkage of the pupil hole upon contraction of an iris-mimicking muscle that had been untethered from its 24-well plate.

**List and description of supplementary CAD files:**

- **Linear_Alignment_StampHolder.STEP:** CAD Model of a single stamp holder used for alignment studies in 24-well plates
- **Linear_Alignment_StampHolder2.STEP:** CAD Model of two stamp holders used for alignment studies connected by a rigid bridge used for alignment studies in Cellvis 24-well plates (Cellvis, P241.5HN). Adjust as needed to fit different 24 well plate formats
- **UpNanoStamp_12-5um.STEP:** CAD Model of the UpNano Resin 12.5 µm stamp used for alignment studies and attached to Linear_Alignment_StampHolder2 as described in the main text

- **UpNanoStamp_25um.STEP:** CAD Model of the UpNano Resin 25 μm stamp used for alignment studies and attached to Linear_Alignment_StampHolder2 as described in the main text
- **UpNanoStamp_125um.STEP:** CAD Model of the UpNano Resin 125 μm stamp used for alignment studies and attached to Linear_Alignment_StampHolder2 as described in the main text
- **UpNanoStamp_flat.STEP:** CAD Model of the UpNano Resin featureless flat stamp used for alignment studies and attached to Linear_Alignment_StampHolder2 as described in the main text
- **Stamp_Holder_Single_Iris.STEP:** CAD Model of a single stamp holder used to make the iris patterned stamps in 24 well plates (Cellvis, P241.5HN) as described in the main text.
- **UpNanoStamp_25um_Iris.STEP:** CAD Model of the UpNano Resin iris stamp used to make the iris patterned stamps and attached to Stamp_Holder_Single_Iris as described in the main text
